# Global monitoring of wildlife mortality through participatory science in near-real time

**DOI:** 10.1101/2025.08.08.669145

**Authors:** Diego Ellis-Soto, Liam U. Taylor, Elizabeth Edson, Avery Hill, Christopher J. Schell, Carl Boettiger, Rebecca F. Johnson

## Abstract

Detection of wildlife mortality events is critical for timely conservation and natural resource management. We present an open-source, web-based decision support tool that queries, aggregates and summarizes participatory science data from iNaturalist to monitor mortality events worldwide. We demonstrate the effectiveness of this approach using four case studies spanning taxonomic, spatial, and temporal scales. In Canada and the United States, high peaks of bird mortality coincided with zoonotic risk during avian influenza outbreaks. Across Latin America, we detected 75 mortality events of critically endangered species. In California, recorded mammal mortality was associated with human infrastructure, including proximity to roads, and to a lesser extent, the human footprint. Mortality of pumas (*Puma concolor*) was detected across nine countries, highlighting the need for international cooperation to conserve mobile species. Our tool enables resource managers to flag emerging threats and empowers participatory scientists to monitor and integrate mortality records for conservation.

## Introduction

The ongoing biodiversity crisis threatens wildlife populations with negative consequences for ecosystem functioning and human health (Pecl *et al*. 2017). Despite ambitious conservation goals imposed by nations worldwide and an increasing number of protected areas, wildlife population declines and extinction risk are ongoing (Runge *et al*. 2015). For example, North American avifauna have experienced a 30% decline in the last 50 years alone with unclear associated mechanistic links (Rosenberg *et al*. 2019). Therefore, in planning conservation interventions and wildlife management, it becomes imperative to understand bottom-up information on population dynamics related to when, where, and how animals die.

In recent decades, advances in biodiversity sensors and data transmission technologies have enabled near-real time biodiversity monitoring at global scales (Besson *et al*. 2022). Consequently, early detection of wildlife mortality has gained increasing attention, allowing managers to respond dynamically to short-term forecasts and shifting ecological conditions. Key efforts include estimating vital rates and detecting mortality events (Sergio *et al*. 2019). Timely detection is especially critical during sudden, extreme events such as heat waves, disease outbreaks, or anthropogenic disturbances, as these can trigger large-scale wildlife die-offs and pose substantial management challenges.

Successful mortality detection has been possible when tailored to specific taxonomic groups (Cossey *et al*. 2025) or event-based surveillance (Kelly *et al*. 2021). With the advance of animal telemetry, animals equipped with bio-loggers have offered a closer look at the causes of individual mortality, identifying particular anthropogenic risks such as power lines for large migratory birds (Serratosa *et al*. 2024), roads, as well as identifying the onset of swine fever infections in wild boar (*Sus scrofa*) (Morelle *et al*. 2023) and the survival or death of individual fish (Villegas-Ríos *et al*. 2020). However, telemetry-based monitoring is often limited to a relatively small number of individuals for larger-bodied species, and geographic areas (Ellis-Soto *et al*. 2025). Thus, despite tremendous potential, most efforts to monitor wildlife mortality events are constrained by geographic, taxonomic or temporal scope.

The rapid increase in participatory science (also known as “citizen or community science”) offers a promising angle to support existing efforts that monitor wildlife mortality (Supplementary Figure 1). To date, participatory science platforms provide the majority of biodiversity data collected worldwide (Callaghan *et al*. 2023) in a manner that is cost-effective, taxonomically expansive (Chandler *et al*. 2017), and accessible across public and private lands. We note there are substantial biases in participatory science datasets, especially in terms of social and economic factors (Ellis-Soto *et al*. 2023; Carlen *et al*. 2024; Chapman *et al*. 2024). Despite these biases, participatory science offers a complementary method to monitor wildlife and identify particular anthropogenic and environmental causes of mortality. For example, iNaturalist, one of the most widely used participatory science platforms, is employed to assess road mortality for diverse taxa (e.g. herpetofauna and mammals (Schwartz *et al*. 2020; Shin *et al*. 2022)) and pinpoint the timing and spatial extent of die-offs in migratory birds (Taylor *et al*. 2024).

Here, we introduce an open-source web-based decision support system to monitor wildlife mortality using crowdsourced iNaturalist records. The records are global in scope and updated daily. We describe the workflow architecture and demonstrate its use with four case studies spanning geographic scales and taxonomic groups. First, we detect mass mortality events associated with H5N1 avian influenza across Canada and the United States, expanding on recent work (Taylor *et al*. 2024). Second, we examine spatial patterns of mortality for large mammals across California, USA. Third, we identify records for species of conservation concern across Latin America. Finally, we showcase the application’s capacity to track mortality in a wide-ranging predator across the Americas, the mountain lion (Puma concolor). Taken together, these case studies highlight the potential for a wide range of conservation and management applications across taxonomic, temporal and geographic scales.

## Materials and Methods

### Mortality records in iNaturalist

iNaturalist is one of the world’s fastest growing and widely used participatory science platforms, with over 265 million observations from over 3.8 million users as of the 8^th^ of August 2025. The site allows users to identify species and upload photographs or audio recordings of organisms along with essential metadata such as location, date, time, and taxonomic identification. iNaturalist plays a vital role in biodiversity data collection, supplying at least 58% of records for all taxa worldwide (https://www.inaturalist.org/blog/76606-thank-you-for-helping-generate-most-gbif-records-for-most-species-since-2020). Data quality is supported through a community-based identification process: for an observation to reach research-grade status, it must have a valid date and location, and at least two users must agree on its species-level identification—often with the observer contributing one of these validations. If consensus is not achieved, the record remains unconfirmed. This may particularly be the case for species that are particularly challenging to identify, or if there are fewer experts in a given taxon, at a given locality, or some times of year attracting less observational effort (Bennett *et al*. 2024).

This robust model enables iNaturalist data to be effectively leveraged for scientific research and wildlife management applications in near-real time worldwide. In contrast to most other participatory science platforms, iNaturalist users can note whether animals are ‘dead’ or ‘alive,’ allowing the public to document mortality events. Another benefit of iNaturalist is that users can join custom projects. This supports the creation of specific local initiatives, fostering and facilitating the sharing of expertise among data collectors and enhancing biodiversity monitoring. These projects help researchers and conservationists better understand the context under which a biodiversity record is generated. One example includes bird strike collision projects which track avian mortality caused by window strikes (e.g. https://www.inaturalist.org/projects/yale-and-new-haven-bird-window-collisions). Place-based events like bioblitzes are also supported, notably the City Nature Challenge, which includes hundreds of city projects. Other local examples include state wide efforts including Roadkill South Africa (https://uk.inaturalist.org/projects/roadkill-s-afr), the Pacific Newt Roadkill (https://www.inaturalist.org/projects/pacific-newt-roadkill-main-project-lexington-reservoir) and beach stranding mortality observation projects in south Australia (https://www.inaturalist.org/projects/sa-marine-mortality-events-2025).

### Deployment of a Shiny Application through Huggingface

Our methodological workflow to identify mortality events consists of a series of queries that pull data directly from the iNaturalist API (https://api.inaturalist.org/v1/). We provide a Shiny Application allowing users to query records based on taxonomic scope, temporal scale and spatial extent. Spatial extent is either defined by providing a specific location via text which will return a bounding box around a place of interest (e.g. Uruguay, or California) or based on a user defined bounding box through a Leaflet map hosted layer (Fig. 1). Our Shiny Application is hosted through a HuggingFace web application (https://huggingface.co/spaces/diegoellissoto/iNaturalist_mortality_detector) and necessary R packages and dependencies are embedded on a Docker Container (Boettiger 2015). We provide documentation on the necessary software and package dependencies employed, and our source code is also available through the Huggingface spaces repository. Given the iNaturalist API’s 240-record limit per query, we recommend users to select smaller spatial extents for near-real time analyses or use our precompiled archive for broader spatiotemporal coverage. To prevent overloading the server, we included a 1.4-second pause between each weekly query for each selected year. To address scalable and large-scale analyses, we compiled an archive of iNaturalist mortality records from the 1st of January 2010 to the 15th of April 2025. This dataset reduces download time and processing latency for retrospective studies and enables exploration of long-term trends without relying on API. We plan to update this archive periodically. Once a user defines a query, our application displays mortality records in space, highlights commonly recorded species, allows data download, and lets users explore individual records through an interactive table with images for verification. To provide users with more local context, we incorporate summary statistics including average daily records, total records, and temporal peaks for the selected area and period of time (Figure 1).

**Figure 1.**
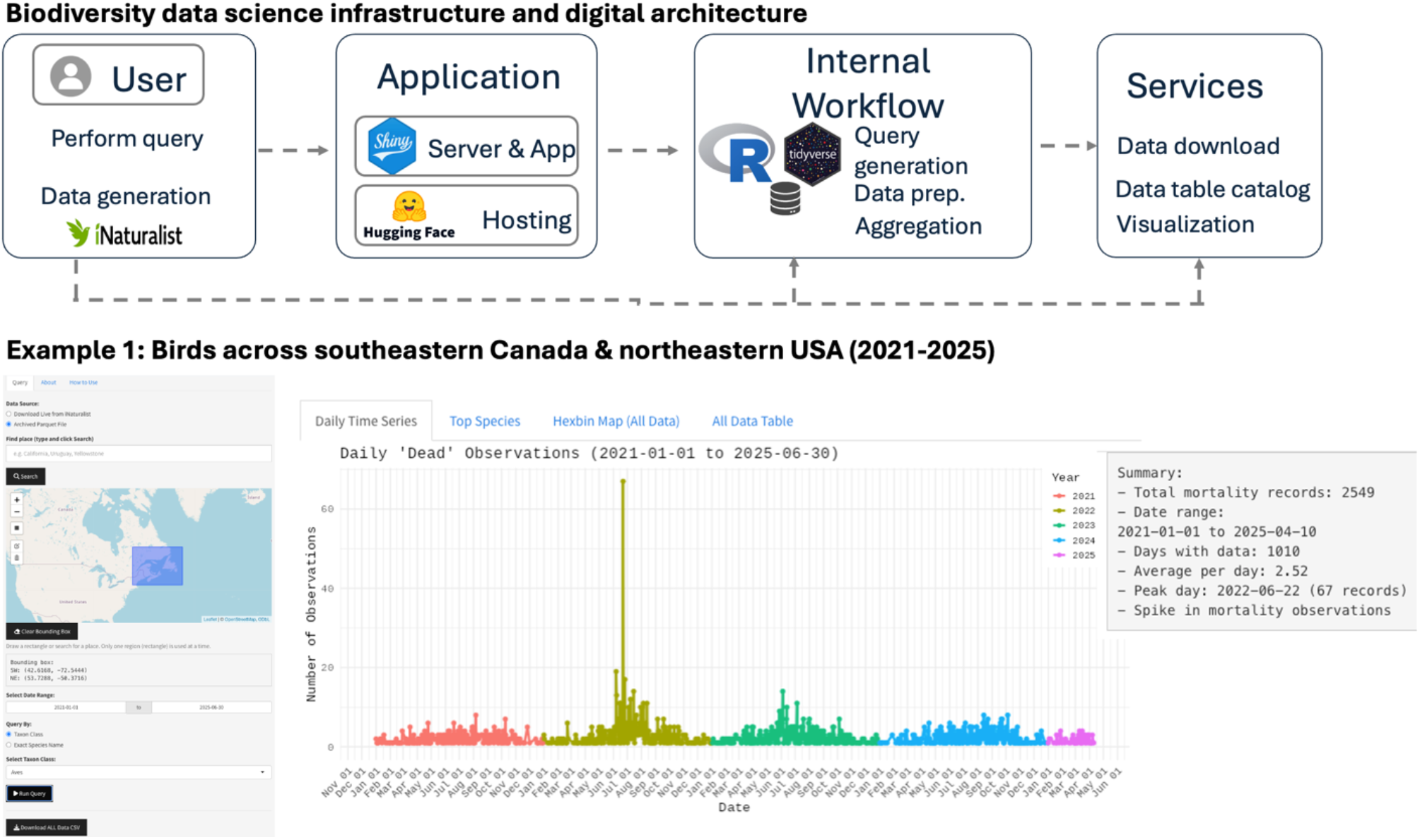
(Top panel) Schematic of the web application for near-real time monitoring of mortality events, integrating querying to the iNaturalist API, a Shiny-based server and app, and hosting via Hugging face. (Bottom panel) Case study demonstrating potential tracking of zoonotic outbreaks in birds across southeastern Canada and northeastern United States. The Hugging Face Application allows users to select a geographic area through manually drawing a bounding box on a map via a leaflet rendered interface. We exemplify how this application aggregates and visualizes trends in mortality detection over time with color coded years. In the highlighted example, a pronounced peak of mass mortality is identified in July 2022, overlapping with the ongoing bird flu H1N5 outbreak of that time.

### Case studies

This series of case studies illustrates the potential of our web-based decision support tools for querying mortality records of specific taxa across geographic regions. First, building on previous monitoring of a bird flu outbreak —integrating iNaturalist data with government-based H1N5 surveillance (Taylor *et al*. 2024)—we analyzed patterns of avian mortality across southeast Canada and northeast United States in 2021 and 2025. Second, we annotated mammalian mortality records across California with anthropogenic variables. Specifically, the human footprint index at 1 km^2^ (Theobald *et al*. 2020), distance to the nearest road, road type using the tigris package in R version 4.4.1. (Walker 2016). This approach both identifies hotspots of data collection and reveals whether certain road types are associated with more mortality records and distinct spatiotemporal patterns in user contributions. We downloaded a total of 8,975 records on the 15^th^ of February 2025, for a total of 285 unique species.

Third, we estimated wildlife mortality across Latin America, particularly for endangered species. This was done to exemplify analysis encompassing diverse ecoregions, social, cultural, and economic human context, and multiple threatened species of conservation concern. We filtered 2023 mortality iNaturalist observations of class Animalia from Latin America to identify IUCN-listed endangered species flagged as deceased. This yielded a targeted list of taxa warranting urgent conservation attention. We also performed this life query on the 15^th^ of February 2025. Finally, we showcased single species mortality of mountain lions (*Puma concolor*) across large spatial extents across the Americas to highlight mortality events of an iconic large predator between 2020-2025. This query was performed on the 20^th^ of June 2025.

## Results

### Bird monitoring revealed spikes in mass mortality associated with H1N5 avian influenza outbreaks

In our first test case, iNaturalist data from the northeastern United States and southeastern Canada successfully captured peaks in bird mortality (defined as two standard deviations above the mean) that aligned temporally with reported outbreaks of avian influenza. These spikes were notable both in their timing—often coinciding with public reports of H1N5 bird flu incidence—and the magnitude of mortalities reported. A major advantage of iNaturalist is that individual user submissions of mortality include photographic evidence, enabling quick community verification. These patterns demonstrate how participatory science can complement official surveillance efforts by identifying potential mass mortality events, thereby providing near-real time information for resource managers and public health officials (*sensu* Taylor et al. 2024).

### Mammal mortality across California

In California, iNaturalist mammal mortality records were concentrated near road networks (>95% of observations occurred within 1,000 m of a road), underscoring the ongoing threat of vehicular collisions to wildlife. Furthermore, mortality records showed a bimodal distribution across the human modification index, with peaks at high and intermediate levels of urbanization (Figure 2). We provide additional information for this analysis on Supplementary Material 1.

**Figure 2.**
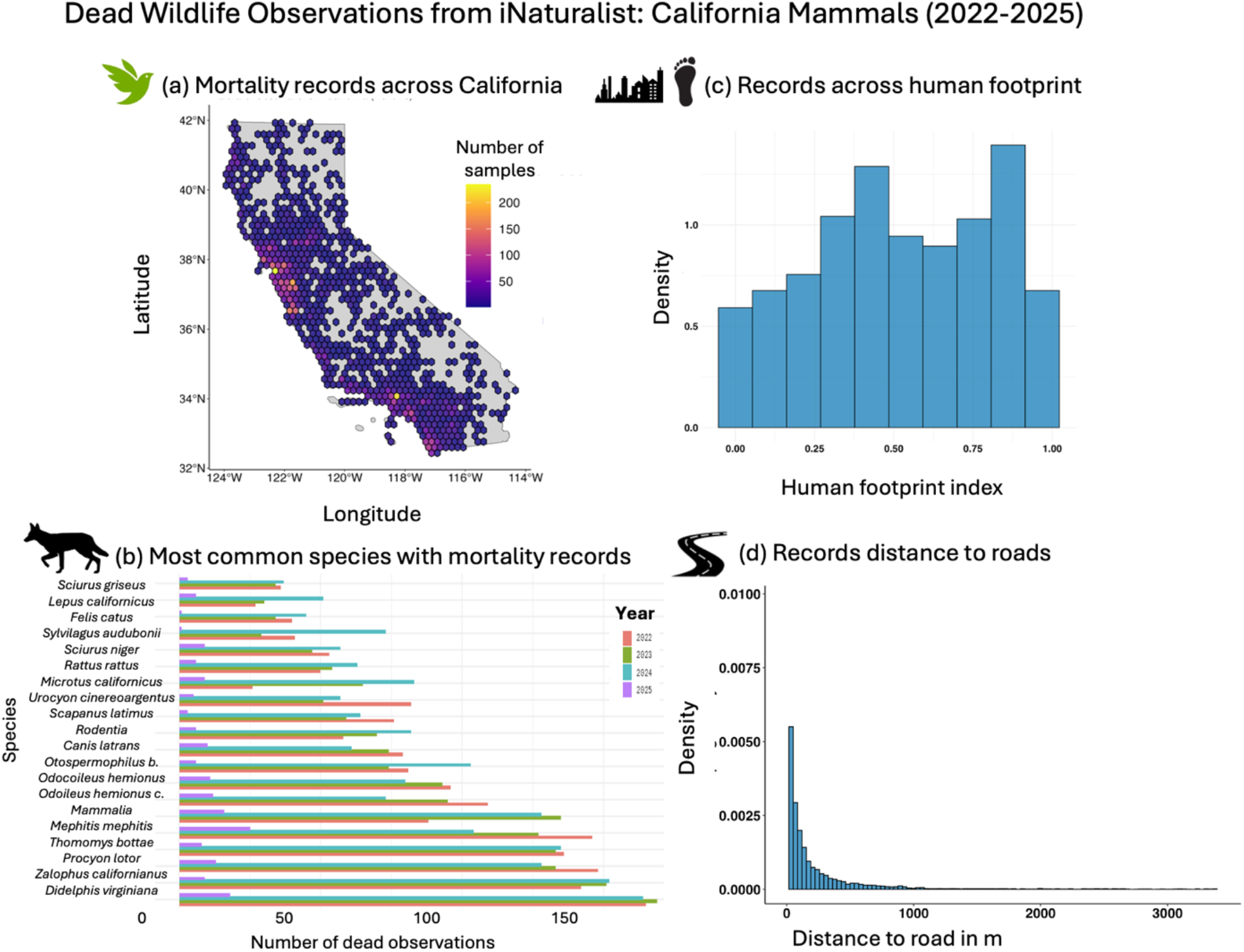
Mammal mortality events across various road types in California, United States. **(a)** Spatial distribution of iNaturalist-recorded mammal mortality events. **(b)** Twenty most common mammal species with mortality records. **(c)** Distribution of records across the Human Footprint Index. **(d)** Distance to nearest road for mammal mortality records across California.

### Detection of endangered species across Latin America

A total of 3,397 mortality observation records were identified across Latin America. Of those, participatory scientists identified a total of 75 observations that came from 11 critically endangered species, 49 from endangered, 48 from near threatened, and 218 observations from vulnerable species after IUCN status. Such information has the potential to identify broader locations where conservation interventions may be urgently needed to protect threatened species.

### The political biogeography of Puma mortality monitoring

We identified 94 deaths from mountain lions from 2020-2025 across a total of nine countries and political boundaries connecting North America, Central America and South America. This included 9 records in Mexico, 18 records in Argentina, 52 in the United States, 4 records in Brazil, 3 in Canada, 4 in Chile, 1 in Colombia, 2 in Costa Rica and 1 in French Guiana (Figure 3).

**Figure 3.**
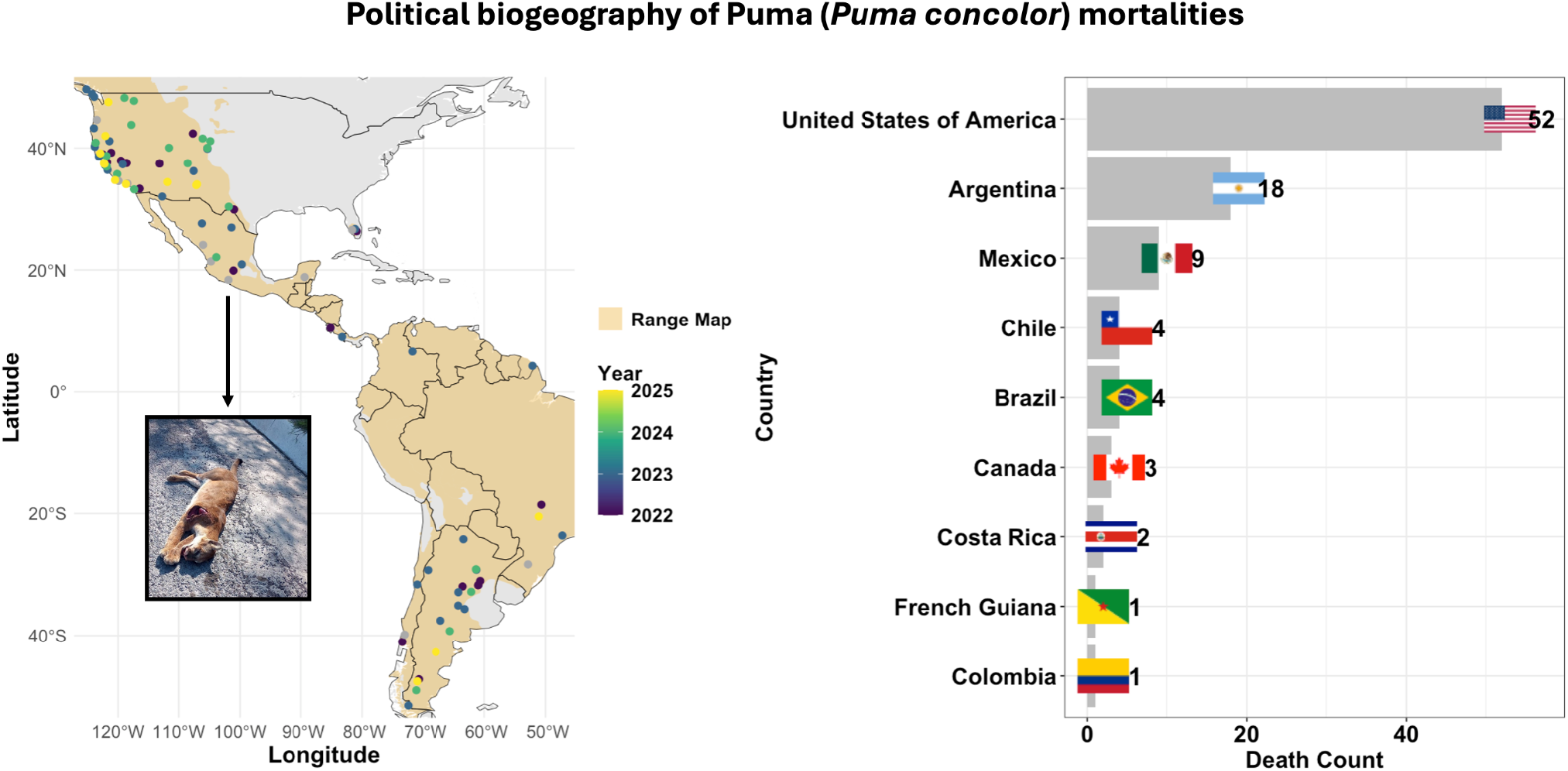
(Left panel) Mortality events of Mountain Lions (*Puma concolor*) across the Americas, spanning a large broad, biogeographic and jurisdictional gradient across nine countries. Species range map was obtained on the 20th of June 2025 from The IUCN Red List of Threatened Species 2015: e.T18868A97216466. https://dx.doi.org/10.2305/IUCN.UK.2015-4.RLTS.T18868A97216466.en. Photograph: deceased puma in Mexico by iNaturalist user record https://www.inaturalist.org/observations/105943003). (Right panel) Bar plot of mortality records across American countries.

## Discussion

Our findings underscore how participatory science can enhance early warning systems for wildlife management and public health by enabling near-real time detection of mortality events across diverse taxa. Our case studies illustrate how the dynamic filtering by taxonomic groups enables stakeholders to prioritize interventions for species or regions of interest. By leveraging iNaturalist’s global reach and daily influx of detailed observations, managers and researchers could quickly identify and respond to emerging threats—such as mass mortalities associated with extreme weather, disease outbreaks, or anthropogenic hazards. Mortality data could further be integrated with ancillary information (e.g., vehicular traffic, road networks, building types, zoonotic surveillance from wastewater, timing of bird migration, early detection of invasive species) to better understand the drivers of mortality and tailor strategies to minimize future risks and mitigate human-wildlife conflicts.

The dual capacity for live, near-real time queries and retrospective analyses of archived iNaturalist records means managers can both flag emerging die-off events— distinguishing whether mortality spikes are localized, cyclical, or widespread—and uncover long-term trends. By leveraging community-based projects (e.g., bird-strike surveys, road mortality projects) with broad-scale mortality observations, the platform offers broad bottom-up decision-support for local conservation efforts, public health officials, resource managers, and biodiversity scientists.

The near-real time nature of our app enables community-driven data curation: users can spot and correct misidentified records in iNaturalist, with updates reflected in future queries—creating a feedback loop that improves data quality over time. Future work could build upon iNaturalist’s existing—but underused—place-and-taxon alert engine and integrate it to our web-based platform to allow managers to receive automated alerts when mortality records for particular taxa or unusually high numbers of observations are reported within a region of interest.

Leveraging participatory science for near-real time monitoring of wildlife mortality requires careful understanding of the limitations and biases of such opportunistic recordings (Taylor *et al*. 2024). Although iNaturalist provides valuable photographic evidence and community-based species identification, not all regions have large user communities, leading to substantial geographic gaps in reporting (Geurts *et al*. 2023). For instance, the higher reported mortality of pumas in the United States relative to other countries may reflect better access to digital infrastructure, and greater use of iNaturalist, rather than true differences in mortality rates.

In less populated or remote areas, a single volunteer might document multiple mortality incidents, whereas in urban centers, numerous users could document the same event multiple times, artificially inflating its apparent magnitude. However, mortality records are perhaps the easiest community-science records for which to spatially and temporally filter out duplicates given that carcasses are less likely to move and should be detectable for limited time periods. Additionally, certain taxa, such as cryptic or nocturnal species, may be underreported because they are harder to detect or identify confidently in the field (Di Cecco *et al*. 2021; Bennett *et al*. 2024). Similarly, common species are also likely to be underreported. For example, dead deer, pigeons or rats are abundant enough that observers might not consider such observations for submission to platforms like iNaturalist.

Further, species dying along heavily trafficked roads may be difficult to note by participatory scientists during transit. To address these issues, integration of participatory science data with additional sources—such as governmental transportation agency records, wildlife rehabilitation centers, or animal control databases—would offer a more comprehensive picture, improving our understanding of the ‘when, where, and how’ metrics of wildlife mortality events. For these reasons, we stress that iNaturalist mortality records alone should be treated with caution when estimating the magnitude of animal mortality. Addressing these limitations through targeted expanded user engagement and partnering with local agencies—especially in underrepresented regions—will enhance the reliability and utility of this forecasting tool.

The timing and frequency of user observations also vary, with some participants being highly active over short periods (e.g., bioblitzes) and others contributing sporadically (Di Cecco *et al*. 2021). Understanding these nuances is essential for interpreting patterns: a spike in observations may reflect a genuine mass mortality event, a concentrated effort by observers, or both. Furthermore, some users may also be uncomfortable posting observations of deceased organisms, leading to underreporting of mortality events. Moreover, in the iNaturalist mobile app, marking an organism as alive or dead requires an extra step after the observation is created via the web interface, which users may easily overlook. A potential solution could involve a campaign encouraging users to flag both their own and other’s observations as dead to improve mortality data reliability and completeness

### Outlook

We see particular premises in future co-development with wildlife management and public health agencies to integrate opportunistic mortality records, as well as with environmental organizations, iNaturalist user groups, and projects. For public health, this could include integrating migratory bird abundance estimates with monitoring of viral loads through wastewater surveillance (Lee *et al*. 2024). Similarly, close engagement with wildlife agencies tasked with managing species of conservation concern could enhance the spatio-temporal coverage of information available across public and private lands. Future research could further identify key sources of human mobility to inform the design of ecological corridors or reduce bird collisions with specific building types. Government agencies could set up automatic message deliveries when daily runs of mortality detection in an area of interest exceeds a threshold, enabling rapid responses for adaptive management of wildlife. With human-wildlife overlap only increasing (Ma *et al*. 2024), better understanding of where, when and why wildlife is declining will become imperative for conservation efforts and natural resource management.

## Acknowledgements

We would like to express our gratitude to all participatory scientists worldwide leveraging biodiversity tools which have made this mortality monitoring decision support tool possible. D.E-S. acknowledges funding support from the David H. Smith Research Fellowship and the Presidential Postdoctoral Program of the University of California. D.E-S. acknowledges feedback from Grant Higerd-Rusli.

## Author contributions

D.E-S performed data analysis and visualization and led writing of the original draft, with significant feedback from L.U.T. All co-authors wrote, reviewed, and edited the manuscript.

## Data and code availability statement

The codes used to download, visualize, and analyze biodiversity data are available on GitHub (https://github.com/diego-ellis-soto/iNat_mortality_detector). This web-based platform is hosted and available on https://huggingface.co/spaces/diegoellissoto/iNaturalist_mortality_detector. All data sets utilized are publicly available and can be sourced from the provided Hugging Face and Github repository.

---

**Supplementary Figure 1.**
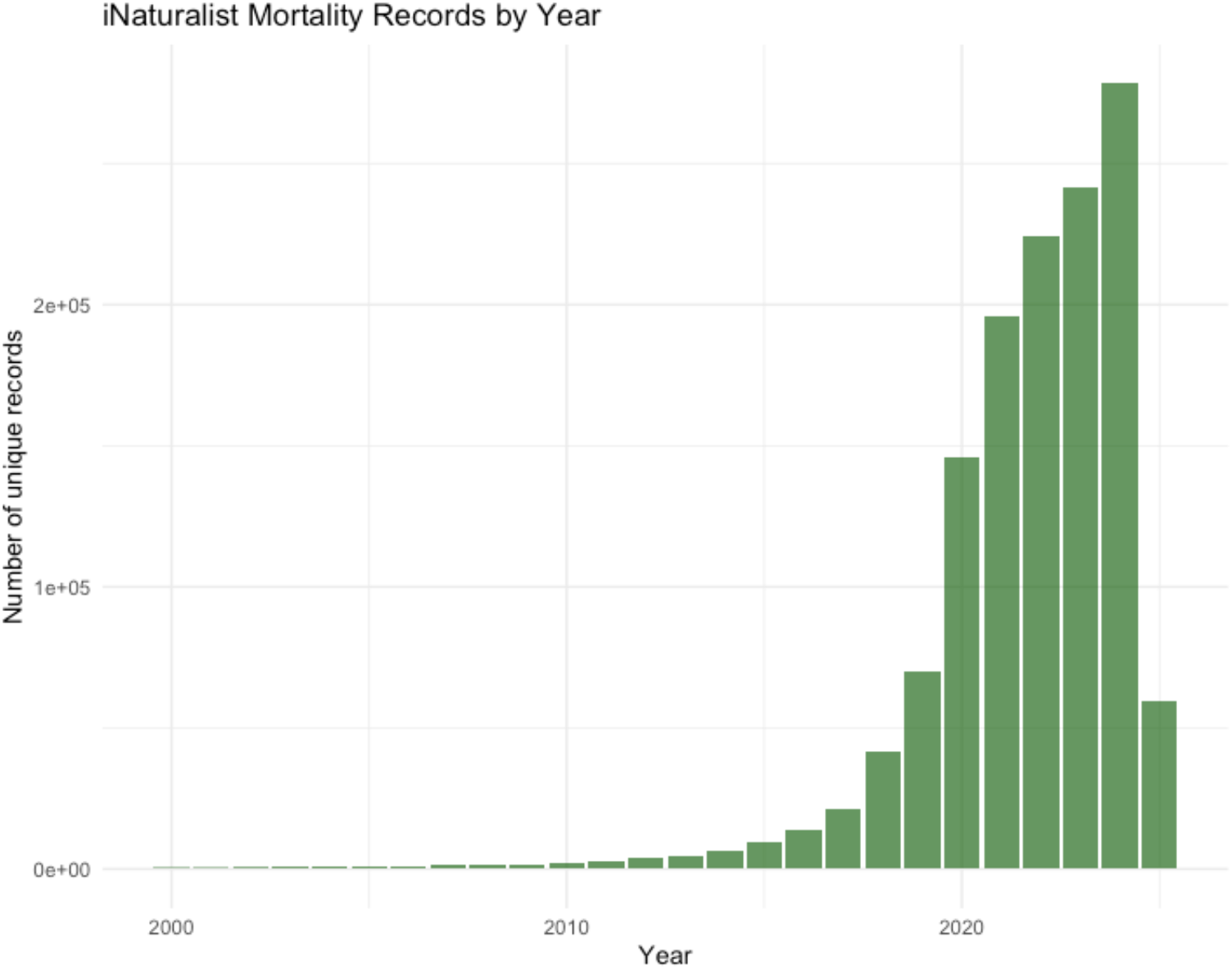
Temporal growth of iNaturalist mortality records since 2000. Note a rapid increase from 2020 onwards.

## Notes

https://huggingface.co/spaces/diegoellissoto/iNaturalist_mortality_detector

### Competing Interest Statement

The authors have declared no competing interest.

