## Supplementary Material 1 for "Global monitoring of wildlife mortality through participatory science in near-real time"

*Assessing mammal mortality across California in relation to distance to nearest road and human footprint*

To understand the association of mammal mortality across California with anthropogenic variables related to infrastructure, such as roads and human modification (Theobald et al. 2016), we performed spatial operations in R version 4.4.1 (R Core Team 2021). Road networks across California were obtained via the *tigris* package (Walker 2016) which allowed us to associate the distance to nearest road for each mortality record, as well as by road type. All data was reprojected to California's Albers projection (EPSG:3310) using the *sf* package for distance calculations and area-based analyses (Pebesma 2018). Distance to nearest road was calculated by identifying, for each mortality point, the nearest road feature using st_nearest_function, extracting the corresponding road geometry, and computing point-to-line distances in meters. We visualized these distances into a density histogram.

The codes used to download, visualize, and analyze biodiversity data are available on GitHub (https://github.com/diego-ellis-soto/iNat_mortality_detector). This web-based platform is hosted and available on https://huggingface.co/spaces/diegoellissoto/iNaturalist_mortality_detector. All data sets utilized are publicly available and can be sourced from the provided Hugging Face and Github repository.

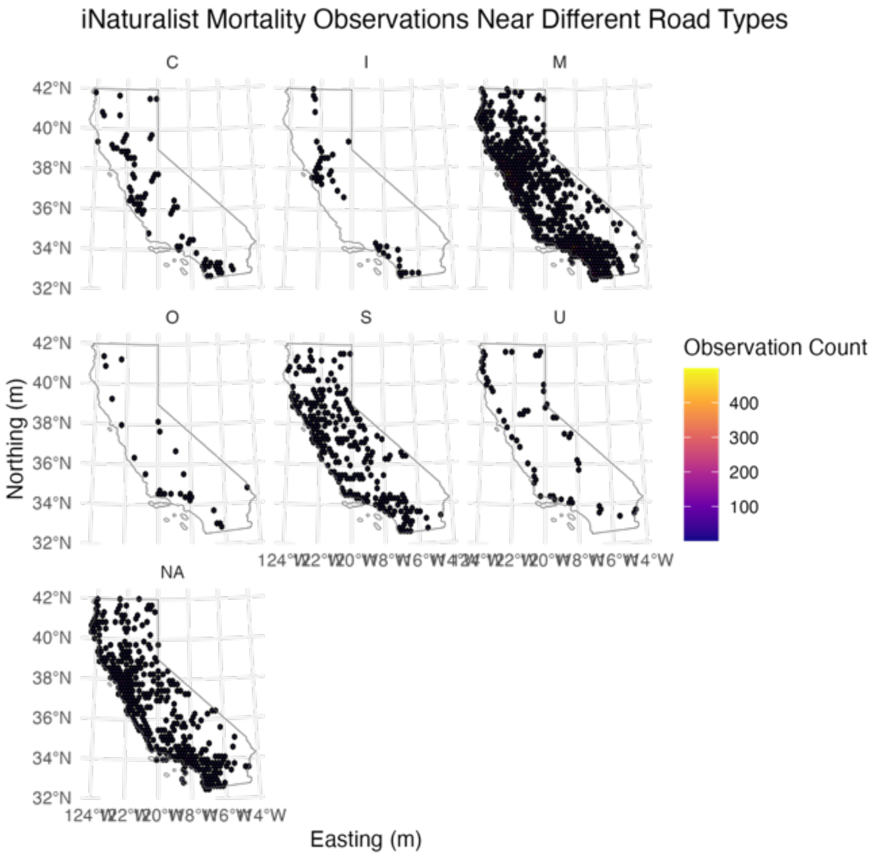


Mortality records for mammals from iNaturalist across different road types across the state of California, USA.
